# Root and canopy traits and adaptability genes explain drought tolerance mechanism in winter wheat

**DOI:** 10.1101/2020.11.04.367904

**Authors:** A.S. Nehe, M. J. Foulkes, I. Ozturk, A. Rasheed, L. York, S.C. Kefauver, F. Ozdemir, A. Morgounov

## Abstract

Bread wheat (*Triticum aestivum* L) is one of main staple food crops worldwide contributing 20% calories in human diet. Drought stress is the main factor limiting yields and threatening to food security, with climate change resulting in more frequent and intense drought. Developing drought-tolerant wheat cultivars is a promising way forward. The use of a holistic approaches that include high-throughput phenotyping and genetic makers in selection could help in accelerating genetic gains. Fifty advanced breeding lines were selected from the CIMMYT Turkey winter wheat breeding program and studied under irrigated and semiarid conditions for two years. High-throughput phenotyping were done for wheat crown root traits using shovelomics techniques and canopy green area and senescence dynamics using vegetation indices (green area using RGB images and Normalized Difference Vegetation Index using spectral reflectance). In addition, genotyping by KASP markers for adaptability genes was done. Overall, under semiarid conditions compared to irrigated conditions yield reduced by 3.09 t ha^−1^ (−46.8%). Significant difference between the treatment and genotype was observed for grain yield and senescence traits. Genotypes responded differently under drought stress. Root traits including shallower nodal root angle under irrigated conditions and root number per shoot under semiarid conditions were associated with increased grain yield. RGB based vegetation index measuring canopy green area at anthesis was more strongly associated with GY than NDVI under drought. Five established functional genes (*PRR73.A1* – flowering time, *TEF-7A* – grain size and weight, *TaCwi.4A* - yield under drought, *Dreb1*-drought tolerance, and *ISBW11.GY.QTL.CANDIDATE-* grain yield) were associated with different drought-tolerance traits in this experiment. We conclude that a combination of high-throughput phenotyping and selection for genetic markers can help to develop drought-tolerant wheat cultivars.

## 1. Introduction

Wheat (*Triticum aestivum* L.) is one of the most important food crops contributing around 20% of calories in the human diet worldwide. However, climate change has resulted in more frequent and intense episodes of drought which affect wheat production (1). Worldwide drought is the single most important factor affecting wheat yields (2) and model-based predictions indicate that there will be 9–12% wheat yield reduction in 21^st^ century without considering the benefits of CO_2_ fertilization and adaptations (3). Developing new drought-tolerant varieties is therefore important to achieve food security in the face of climate change. Identifying drought-tolerance traits for deployment in breeding is therefore crucial. Deeper root systems and cooler canopy temperature are two important traits reported to be responsible for drought tolerance in wheat (4,5). As roots are difficult to study under field conditions, canopy temperature has been applied as an indirect way of assessing the role of the root system as higher water transpiration and uptake is related to a cooler canopy. Canopy stay-green characters have also shown promise to help select drought-tolerance genotypes (6,7).

Wheat root systems consist of seminal roots (up to 6) and crown roots (around 10-15) per plant emerging from basal node of main shoots and tillers (8). These two root systems function together to acquire water and nutrients from the soil (9–11). Distribution of root length density (root length per unit soil volume; RLD) with depth is one of the important traits responsible for water capture in wheat crops (6,9,12). In synthetic wheat-derived lines, yield increase under water stress conditions was associated with increase in root dry weight at depth in NW Mexico (13). Higher allocation of plant assimilates to deeper roots was responsible for a cooler canopy and increase in overall grain yield under drought conditions in synthetic derived material (5). Narrower root angle in the top soil (steeper roots) was associated with higher root density in deeper soil in Australia (13–15).

The high-throughput phenotyping of root system architecture under the field conditions presents a bottleneck in breeding for drought tolerance in wheat (16). Previously, the soil-core break method (17) and ‘shovelomics’ (18) have been used for high-throughput field phenotyping in cereals. Shovelomics, which mainly focuses on crown root phenotyping, involves the excavation of roots in the topsoil and measuring root traits manually or through image analysis. Using this method results of direct measurements and visual scoring in maize showed correlations with root depth for crown root number and angle (18). Shovelomics methods have quantified genetic variation in crown root angle and root length in maize (18–20), barley (21) and durum wheat (22). We used a high-throughput shovelomics technique for phenotyping root crown architecture of the whole root crown in wheat, as developed by (23).

The use of Normalized Difference Vegetation Index (NDVI) spectral reflectance index to study canopy growth and senescence dynamics has been common for decades, but it does have its limitations (24,25). Especially at high values of leaf area index (LAI), NDVI tends to saturate and does not show as strong of a linear association with yield components (26,27). Also, in order to obtain accurate results with NDVI, bright conditions with direct sunlight are required while taking the measurements (in the case of passive sensors). Senescence is a genetically programmed and environmentally influenced process (Thomas and Howarth, 2000) and the stay-green phenotype has shown proven utility to improve yields under drought (Borrell et al., 2000; Verma et al., 2004). NDVI has been used to measure stay green in wheat under drought (7). RGB image-based vegetation indexes have proved to be better associated with grain yield than NDVI under similar circumstances and also are time-saving (27).

Marker-assisted selection is very important component of molecular breeding to develop resilient cultivars by selecting and accumulating favorable alleles. In bread wheat several genes underpinning drought adaptability have been identified and molecular markers have been developed to select favorable alleles. However, the distribution and association of such functional genes like *Derb1, PRR-73, TaCwi-A1* and *TEF-7* is largely unknown in wheat cultivars from the most parts of the world. Transcription factors like *DREB* known to control the expression of several functional genes responsible for plant tolerance to drought, high-salt and cold stress and have been propose to use in plant improvement for stress tolerance (28). Similarly, *TaTEF-7A* is transcript elongation factor gene responsible for number per spike (29) and thousand grain weight under (30). The gene *TaPRR73* was found to be regulation of flowering date and can be used in breeding to develop cultivars adaptable for different geographical areas (31). *TaCwi-A1* gene produce cell wall invertase enzyme manly responsible for sink tissue development and carbon allocation and showed affecting grain weight in wheat (32).

The present study reports associations between nodal root traits measured using the wheat shovelomics techniques along with vegetation indexes in a set of 50 CIMMYT Turkey winter wheat cultivars and advanced lines. The experiment was conducted under irrigated (IR) and semiarid (SA) field conditions in Turkey in two years. The germplasm was also screened for allelic variation of genes previously related to drought adaptability genes using breeder friendly KASP markers. Our aim was to identify genes and traits to use in winter wheat breeding for drought tolerance in Mediterranean region.

## 2. Materials and methods

### 2.1 Experimental design and plot management

Two field experiments were conducted at Bahri Dagdas International Agricultural Research Institute, Konya in 2017–18 and 2018–19. Before sowing the experimental field was fallow. The soil type was a sandy clay. Experiments were conducted in a randomized block, split– plot design, in which two irrigation treatments (IR: drip-irrigated and SA: semiarid/rain-fed) were randomized on main-plots, and 50 CIMMYT winter wheat cultivars and advanced lines including 4 check cultivars were randomized on sub-plots in two replicates. The 50 winter wheat genotypes represent modern germplasm developed by International Winter Wheat Improvement Program and obtained from cooperators in Eastern Europe. They were selected on the basis of their performance under both irrigated and semiarid conditions in advanced cultivar trials. The check cultivars used were Gerek, Katea, Konya and Nacibey (S Table 1). Plots were 7.0 m × 1.2 m with 6 rows 20 cm apart and 450 seeds were sown per square meter. Fertilizers applied were 100 kg ha^−1^ of phosphorus (P) and 39 kg ha^−1^ of nitrogen (as ammonium nitrate) per hectare at the time of planting, and an additional 50 kg ha^−1^ of N at tillering (GS35). Under the irrigated treatment drip-irrigation was given as 50 mm application each time. Irrigation was given twice during the crop growth season at tillering and flowering stage.

### 2.2 Crop measurements: Grain yield and yield components

In 2018, a 1.5 m row bulk sample was hand-harvested by cutting at ground level at physiological maturity (GS89). The fertile shoots (those with an ear) were counted and 5 primary (large ear and stem) fertile shoots were selected for dry matter partitioning analysis. In 2019, around 10-20 shoots were selected for dry matter partitioning analysis from the sample for root measurements (next section). All selected shoots were separated into ears and straw. Dry weight of the ears and the straw was recorded after drying at 80°C for 48 h. The ears of the bulk sample were then hand threshed and grain weighed. All grains were counted by a Contador seed counter (Pfeuffer, Germany) and 1,000 grain weight (TGW) was calculated. From these data the grain DM per fertile shoot, harvest index (HI; grain DM / above-ground DM), fruiting efficiency (grain weight per ear dry weight) and above ground dry matter (AGDM; GY/HI) were calculated. The grain yield was calculated by weighing grain from rest of the plot which was machine-harvested (adjusted to 85% dry weight).

### 2.3 Shovelomics root crown trait measurements

The methodology for assessing root crown traits in both years was similar with some modification. Root crowns were excavated from all sub-plots during late-grain filling. A spade of 25 cm width and 30 cm depth was inserted to 20 cm depth on either side of plants keeping blade parallel to the row. Single sample were taken per plot. The soil was placed into a 10 L bucket filled with water for overnight. Next day root crowns sprayed with low pressure water from a hose to remove remaining soil. Three plant per sample where selected for scanning or image analysis. The number of fertile shoots for each of the three plants was counted. In 2018, root images were acquired and analyzed using WinRHIZO Regular V. 2009c scanner and software (Regent Instruments Inc., Canada). The traits measured were root surface area (cm^2^), root diameter (mm), and root volume (cm^3^). In 2019, images of the roots were taken with a RGB camera (Sony a 6000). A single image per sample was taken with auto setting. Roots were placed over black background to maximum contrast and sample ID and reference scale (white square of 2 cm x 1 cm) was placed on side of the roots as shown in S Fig. 1. Images were analyzed using a modified method from York and Lynch (33). A project for the ObjectJ plugin (https://sils.fnwi.uva.nl/bcb/objectj) for ImageJ (34) was created to allow the angles, numbers, stem diameter and roots diameter to be measured from the plant-root samples (S Fig. 1). The pixel dimensions were converted to physical units using measurements of the known-sized scale in every image. The traits measured were root number per shoot, root diameter (mm), and root angle (°). A polyline was used to measure the crown lengths of the outermost roots and the seminal root length, and the angle was measured for the outermost crown roots at approximately 5 cm depth by measuring the width then later calculating angle using trigonometry and the actual depth measurement to where width was measured. For nodal root number, each nodal root axis was manually annotated, and the count recorded in an output file. The image analysis gave values for the number of pixels corresponding to root diameter and numbers. Using the 2 cm x 1 cm reference square, these pixel values were then converted to the relevant units for each root measurement in excel that also calculated angles as detailed in (33).

### 2.4 NDVI and RGB based vegetation indexes and canopy temperature

In 2018, Normalized Difference Vegetation Index (NDVI) was measured using the handheld active sensor Trimble GreenSeeker spectroradiometer (Trimble Navigation Ltd, USA) to assess the canopy green area starting from booting (GS41). Modified Gompertz curves (Eq. 1) were fitted to the NDVI values against thermal time (base temp. 0°C after anthesis, GS61, Zadoks et al., 1974). The Gompertz (T) parameter was fitted as the thermal time for the NDVI to decrease to 37% NDVI value at GS61. The thermal time (t, measured in °CD) when NDVI values were 90% and 10% of the value at GS61 were taken as the onset of senescence (SenSt) and end of senescence (SenEnd), respectively, whereas the duration from 90% NDVI to 10% NDVI remaining was considered as the senescence duration (SenDu).

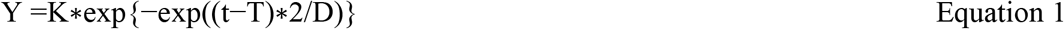

where t is thermal time (base temp. 0°C), D is duration of senescence (SenDu), T is the timing of the inflection point at 37% NDVI value remaining from initial point at GS61, K value is the maximum NDVI at GS61. The senescence parameters were estimated for each sub-plot and then subjected to ANOVA.

In 2019, RGB images and NDVI (GreenSeeker Trimble Navigation Ltd, USA) were taken every two weeks from tillering (GS35) to crop maturity (GS89). RGB image-based vegetation index - green area per meter square (GA m^−2^) was calculated using equation 2 to 6:

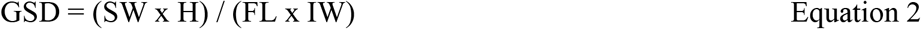

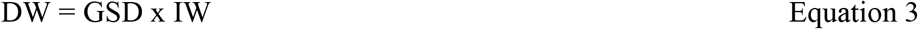

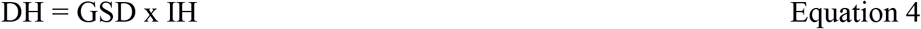

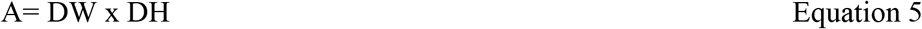

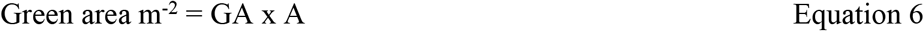

whereas GSD is ground sampling distance (centimeters/pixel), SW is camera sensor width (mm), H is camera height from top of the canopy (m), FL is focal length of camera (mm), DW is width of single image footprint on the ground (m), DH is height of single image footprint on the ground (m), IW is image width (pixels), IH is image height (pixels), A is ground area in image (m^2^) and GA is index value output after image analysis using BreedPix. BreedPix is open source software (35), implemented as part of the open-source CerealScanner plugin (Fernandez-Gallego et al., 2019, https://gitlab.com/sckefauver/cerealscanner) developed for ImageJ software (34). Green Area per meter square at anthesis (GA An) and 2 weeks after anthesis (GA 2W) values are used in this paper. Canopy temperature was measured at anthesis using handheld infrared temperature meter (SBRMART GM320).

### 2.5 Genotyping

DNA was extracted from all genotypes using a modified CTAB method (36). Allele-specific KASP markers for five different loci were used. The primer sequences and amplification conditions of each gene are described in Khalid et al. (2019). The detailed genotyping procedures have been described elsewhere (37,38). Briefly, two allele-specific primers carrying standard FAM tail (5’-GAAGGTGACCAAGTTCATGCT-3’) and HEX tail (5’-GAAGGTCGGAGTCAACGGATT-3’), with targeting SNP at the 3’end, and a common reverse primer were synthesized. The primer mixture included 46 μl ddH_2_O, 30 μl common primer (100 μM) and 12 μl of each tailed primer (100 μM). Assays were tested in 384-well format and set up as 5 μl reaction [2.2 μl DNA (10–20 ng/μl), 2.5 μl of 2XKASP master mixture and 0.056 μl primer mixture]. PCR cycling was performed using the following protocol: hot start at 95°C for 15 min, followed by ten touchdown cycles (95°C for 20 s; touchdown 65°C–1°C per cycle 25 s) further followed by 30 cycles of amplification (95°C for 10 s; 57°C for 60 s). The extension step is unnecessary as amplicon is less than 120 bp. The plate was read in BioTek H1 system and data analysis was performed manually using Klustercaller software (version 2.22.0.5; LGC Hoddesdon, United Kingdom).

### 2.6 Marker-traits association analysis

In this paper we presented results of five key KASP markers out of 150 for their association with phenotypes (Table 5). These markers are regularly used in marker-assisted selection in CIMMYT’s wheat breeding program. As some of the traits that we measured differed between years, the marker-traits associations (MTAs) were identified separately for each year. KASP markers for which one of the alleles was represented at relatively higher frequency (>80%) than other alleles were not considered for MTA. MTA analysis was done in R using liner model (lm) function (Eq. 7) to see the significant effect of the marker on traits by comparing the mean. BLUEs (best linear unbiased estimator) were calculated using Meta-R for randomized block design for all the traits for individual years. BLUE values were used to see the significant effect of marker on traits using following liner model.

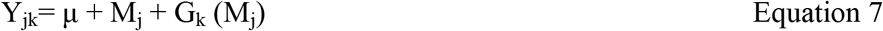

Y is phenotyping value, μ is mean of the population, M is mean effect of j^th^ marker, G_k_(M_j_) genotype within marker variance (error variance).

### 2.7 Statistics

In both years, GenStat 19th edition (VSN International, Hemel Hempstead, UK) was used for carrying out analysis of variance (ANOVA) of traits applying a split-plot design with replications regarded as random effects and genotypes as a fixed effect, and the least significant difference (LSD) test was used to compare the means between specific treatments. A cross-year ANOVA was applied to analyze irrigation treatments and genotypes effects across years and the interaction with year, assuming irrigation treatments and genotypes were fixed effects and replicates and year were random effects. Pearson’s correlation coefficient (r) and linear regressions coefficient (R^2^) were calculated to quantify associations between traits for individual year and cross year means using GenStat. Principal component analyses were done to produce biplots using R software package “factoextra.”

## 3. Results

### 3.1 Drought effects on plant growth

Averaging across the 50 genotypes, the drought/semiarid (SA) conditions reduced the grain yield (GY) compared to irrigated (IR) conditions by 2.67 t ha^−1^ (−50.1%) in 2018 and 3.51 t ha^−1^ (−44.6%) in 2019 (P < 0.001; Table 2) with an average reduction over two years of 3.09 t ha^−1^ (−46.8%, P=0.01, Fig. 1). The cross-year ANOVA showed a significant Year x Genotype (Y x G) interaction (P<0.001, Table 2). Relative loss in GY under SA conditions ranged from −36.1% (genotype code no. 32) compared to −58.5% (genotype 9). The three-way interaction of Y x T x G (Year x Treatment x Genotype) was not significant.

**Table 1.** Environmental conditions during two field crop growing seasons (2018 and 2019) at experimental site Konya, Turkey. Monthly temperature means, (minimum, maximum) and monthly rainfall.

|  | 2017-18 |  | 2018-19 |  |
| --- | --- | --- | --- | --- |
|  | Temperature (°C)<br>Mean (Min, Max) | Rainfall<br>(mm) | Temperature (°C)<br>Mean (Min, Max) | Rainfall<br>(mm) |
| Nov | 12.2 (-0.36, 12.5) | 70.0 | 6.52 (0.92, 12.1) | 27.4 |
| Dec | 7.60 (-2.02, 9.6) | 18.6 | 3.29 (-0.47, 7.05) | 63.4 |
| Jan | 3.22 (-2.66, 5.88) | 4.60 | 0.75 (-3.92, 5.44) | 66.6 |
| Feb | 11.9 (-0.35, 12.2) | 0.20 | 4.25 (-0.99, 9.49) | 31.6 |
| Mar | 19.7 (1.86, 17.7) | 36.0 | 6.24 (-0.78, 13.3) | 20.8 |
| Apr | 26.8 (4.95, 21.8) | 14.4 | 9.59 (2.63, 16.5) | 32.0 |
| May | 35.0 (9.45, 25.5) | 72.2 | 17.12 (7.98, 26.2) | 10.2 |
| Jun | 41.7 (12.3, 29.3) | 38.8 | 21.23 (13.4, 29.0) | 45.6 |
| Jul | 48.8 (16.6, 32.2) | 20.4 | 22.76 (15.3, 30.1) | 7.60 |
| Total |  | 275.2 |  | 305.2 |

**Table 2.** Yield components traits for 50 CIMMYT winter wheat genotypes for 2018, 2019 and cross-year means (CY). Grain yield (GY t ha^−1^), above ground dry matter (AGDM t ha^−1^), harvest index (HI), plant height (PH cm), days to heading (HD), thousand grain weight (TGW), and fruiting efficiency (FE grains g^−1^).

| Genotype | GY t ha<sup>-1</sup> |  | BM t ha<sup>-1</sup> |  | HI |  | PH cm |  | HD |  | TKW g |  | FE grains g<sup>-1</sup> |  |
| --- | --- | --- | --- | --- | --- | --- | --- | --- | --- | --- | --- | --- | --- | --- |
|  | IR | SA | IR | SA | IR | SA | IR | SA | IR | SA | IR | SA | IR | SA |
| Mean 2018 | 5.33 | 2.67 | 12.9 | 5.97 | 0.42 | 0.30 | 86.5 | 52.4 | 122 | 121 | 37.3 | 35.8 | 19.4 | 15.4 |
| Min 2018 | 4.22 | 1.61 | 9.9 | 3.57 | 0.30 | 0.23 | 70.5 | 39.0 | 118 | 118 | 23.4 | 28.2 | 14.3 | 10.4 |
| Max 2018 | 6.56 | 3.76 | 17.5 | 8.88 | 0.48 | 0.36 | 98.0 | 63.5 | 129 | 128 | 50.5 | 44.2 | 26.4 | 19.6 |
| Mean 2019 | 7.94 | 4.37 | 26.6 | 18.0 | 0.43 | 0.25 | 84.9 | 63.4 | 194 | 191 | 33.5 | 30.7 | 21.9 | 15.3 |
| Min 2019 | 6.16 | 2.55 | 21.5 | 13.0 | 0.27 | 0.17 | 73.0 | 50.0 | 186 | 186 | 25.1 | 24.2 | 17.1 | 11.1 |
| Max 2019 | 9.62 | 5.77 | 35.0 | 24.0 | 0.52 | 0.31 | 96.0 | 74.0 | 201 | 196 | 39.6 | 38.5 | 26.8 | 18.6 |
| **CY Mean** | **6.61** | **3.51** | **19.7** | **12.1** | **0.36** | **0.34** | **83.7** | **57.3** | **182** | **180** | **36.6** | **32.1** | **17.4** | **18.6** |
| CY Min | 5.25 | 2.48 | 16.8 | 9.2 | 0.28 | 0.23 | 72.1 | 46.8 | 176 | 176 | 26.6 | 25.7 | 14.5 | 14.8 |
| CY Max | 7.81 | 4.37 | 23.8 | 15.9 | 0.41 | 0.40 | 94.6 | 68.2 | 188 | 186 | 45.7 | 37.7 | 21.4 | 22.2 |
|  | LSD |  | LSD |  | LSD |  | LSD |  | LSD |  | LSD |  | LSD |  |
| G (Genotype) | 0.70 | *** | 2.95 | ** | 0.04 | *** | 4.20 | *** | 1.40 | *** | 2.81 | *** | 1.86 | *** |
| T (Treatment) | 1.47 | ** | 2.9 | ** | 0.01 | * | 7.60 | ** | 0.50 | *** | 3.84 | * | 1.80 |  |
| Y (Year) | 0.60 | *** | 2.24 | *** | 0.05 | ** | 5.50 | * | 1.50 | *** | 1.27 | * | 1.25 | *** |
| T*G | 1.27 |  | 4.35 |  | 0.06 |  | 7.10 | * | 1.90 | ** | 4.38 | * | 2.74 | ** |
| Y*G | 1.02 | *** | 4.25 |  | 0.06 | * | 6.50 |  | 2.10 | *** | 3.98 | *** | 2.67 | *** |

Significance levels displayed as ns > .05, \* < .05 > .01, \*\* < .01, \*\*\* < 0.001.

**Figure 1:**
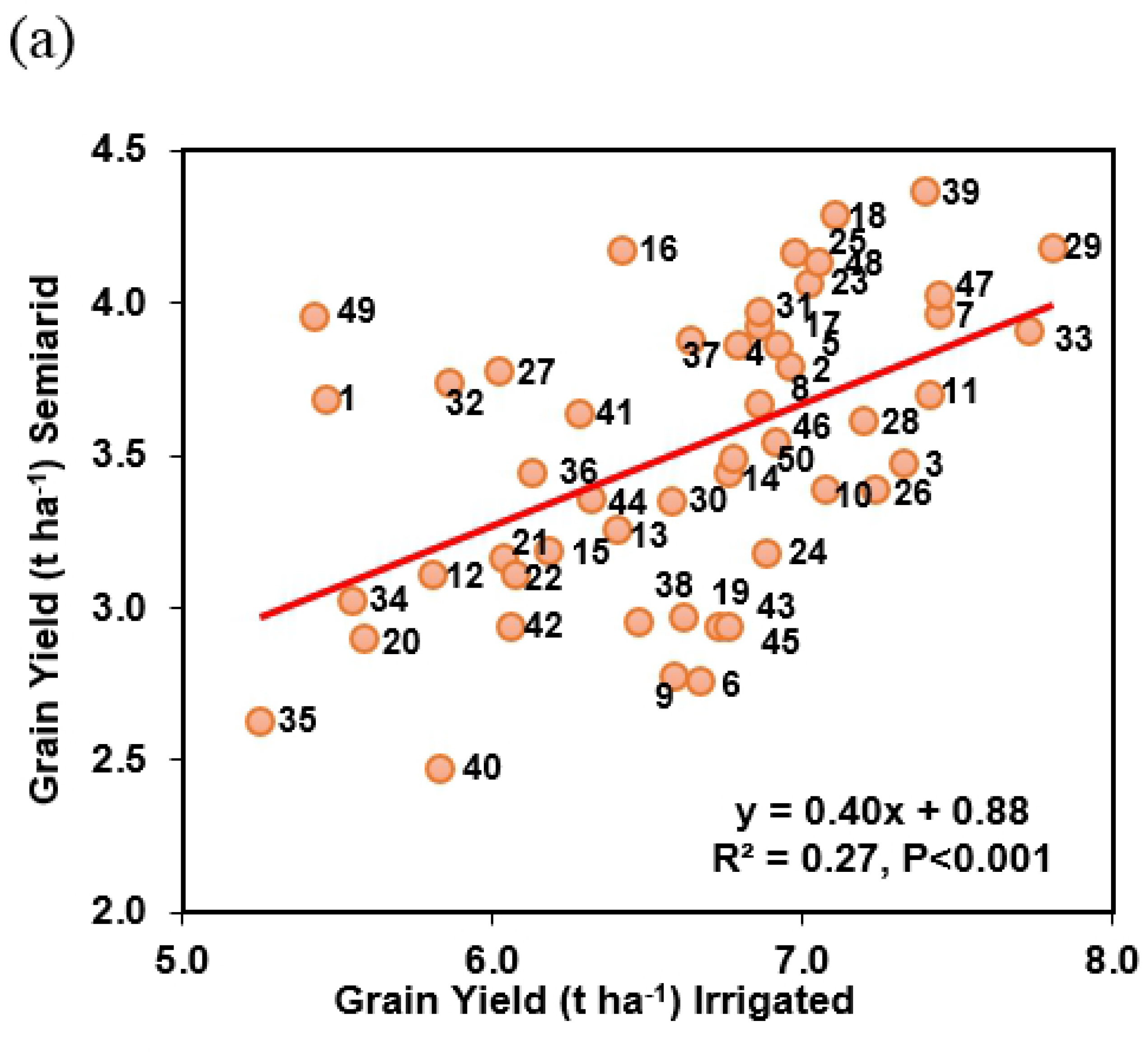
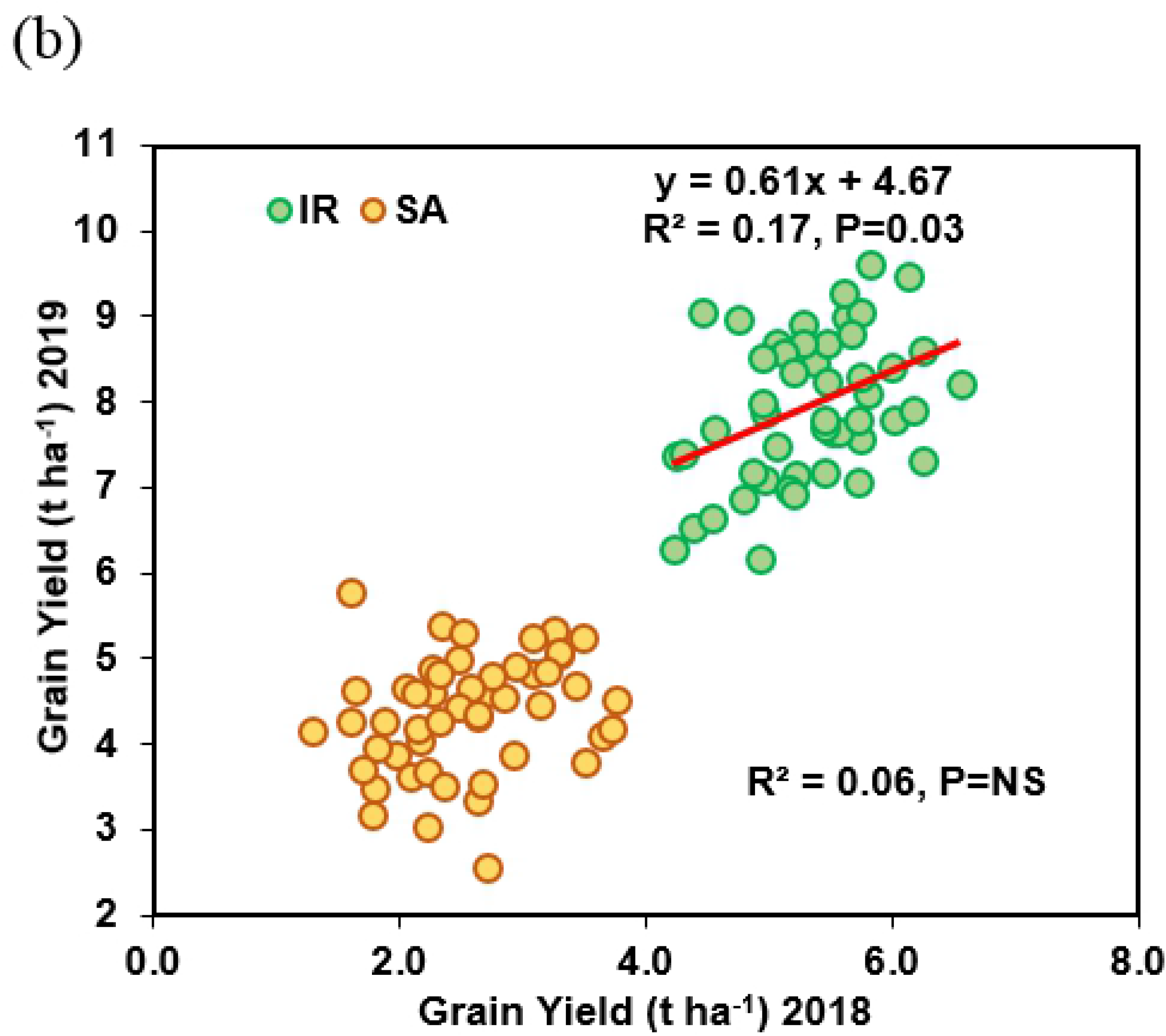
Linear regression among the 50 winter wheat genotypes for (a) grain yield in semiarid on GY in irrigated conditions (mean across the years) and (b) GY in 2018 on GY in 2019 under Irrigated (IR) and semiarid (SA) conditions.

Regression analysis showed a positive association amongst the genotypes for GY between IR and SA conditions (R^2^=0.27, P<0.001, Fig. 1 a). Nevertheless, some genotypes changed rankings markedly; for example: genotype 33 dropped from 2^nd^ highest under IR to 13^th^ highest under SA conditions; 16 and 25 also changed rankings with relatively higher rankings under SA conditions than IR conditions (Table S2). GY also showed positive association with AGDM (IR: R^2^=0.21, P<0.001 and SA: R^2^=0.22., P<0.001) and NDVI at anthesis (IR: R^2^=0.18, P=0.01 and SA: R^2^=0.23., P<0.001) under both IR and SA conditions (Fig. 2 a and c).

**Figure 2:**
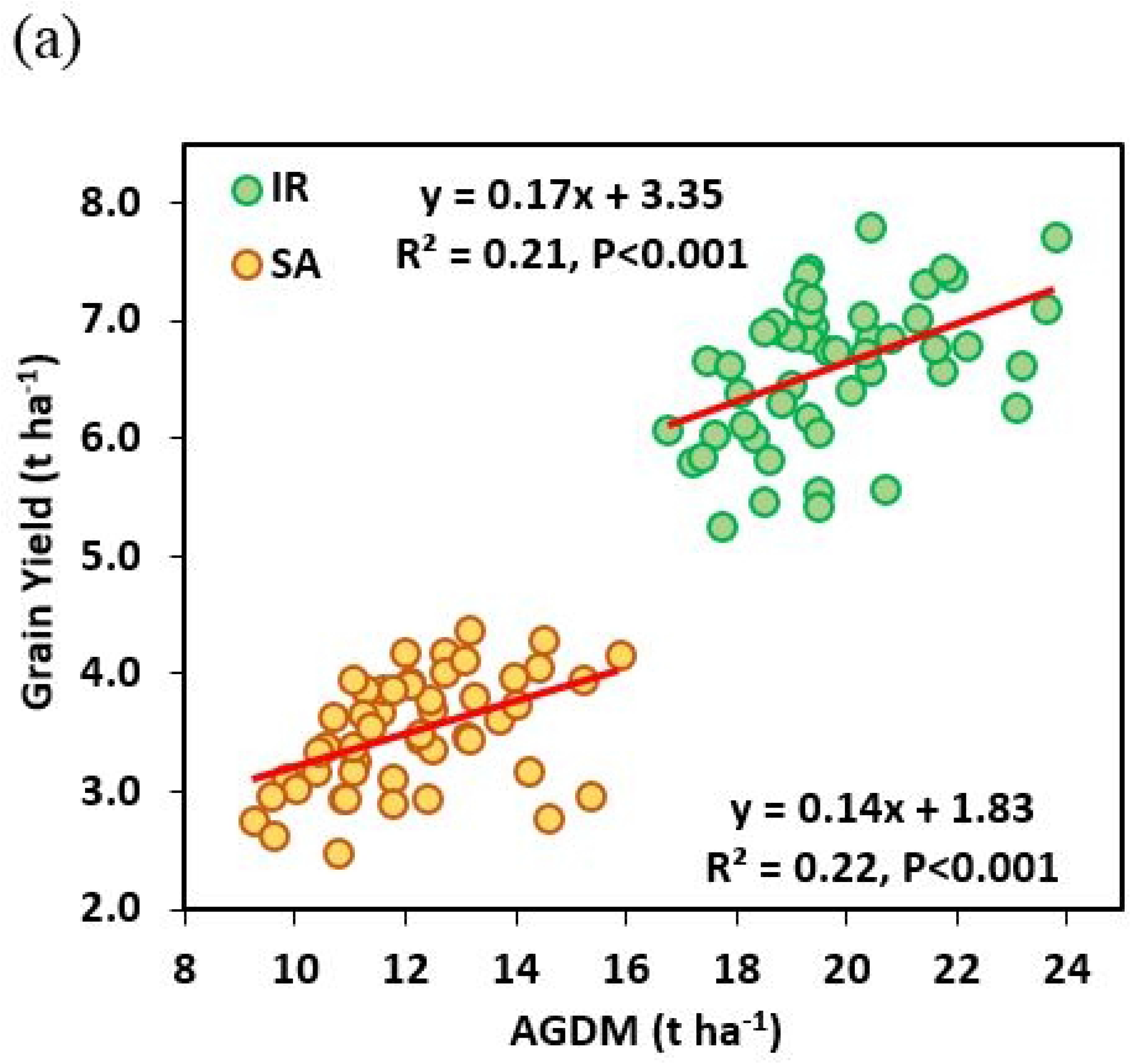
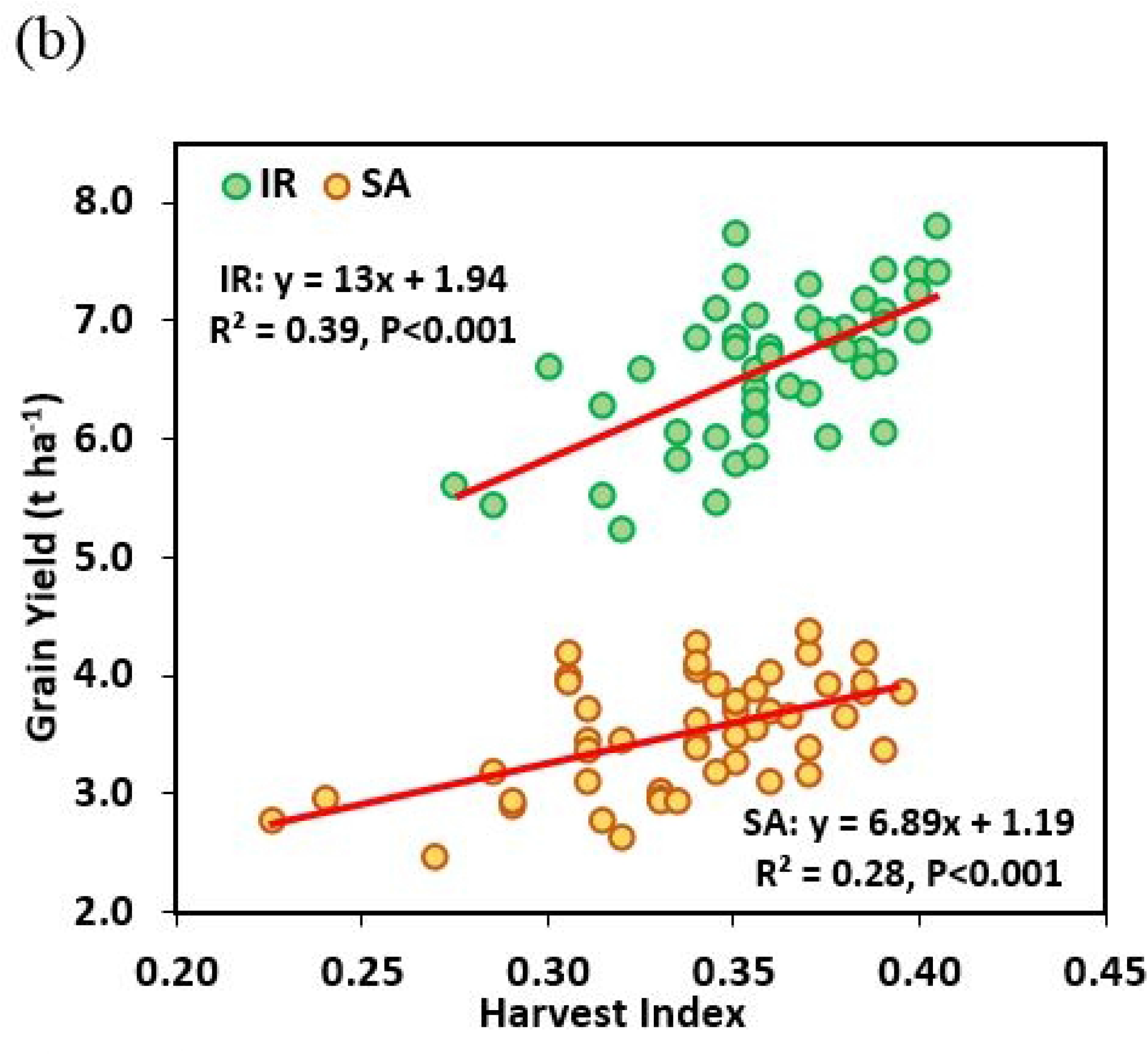
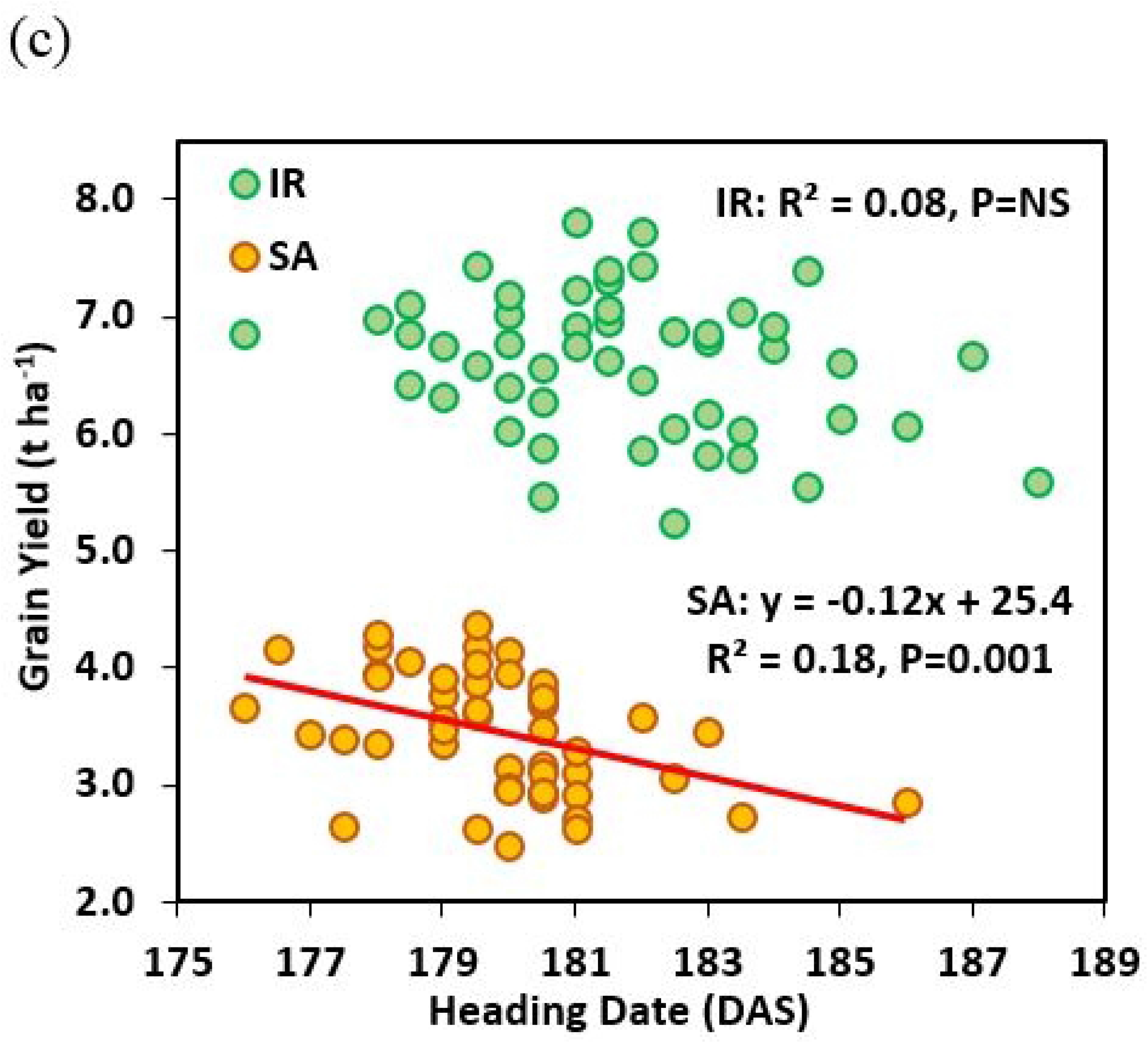
Linear regression amongst 50 wheat genotypes between GY and (a) above ground dry matter (AGDM) (b) harvest index and (c) Heading date (DAS) under irrigated (IR) and semiarid (SA) conditions (mean of 2018 and 2019).

For harvest traits, averaging across years, overall the AGDM was the component affected most by the semiarid conditions reducing from 19.7 to 12.1 t ha^−1^ (−38.6%, P=0.01, Table 2); whereas thousand grain weight (TGW) reduced from 36.6 to 32.1 (−12.2%, P=0.03, Table 2). For AGDM, T x G interaction was not significant whereas for TGW it was (P=0.03). Reduction in grains per ear under SA conditions ranged from −0.5% (genotype 16) to −49.0% (genotype 9). Variation in response to drought for TGW was from −0.9 to −25.7%. Heading date (HD) was advanced by two days in SA conditions compared to IR (P=0.004). There was a negative association between GY and HD amongst cultivars under SA conditions (R^2^=0.18, P=0.002), but no association under IR conditions (Fig. 2 c). Overall drought reduced plant height by 31.6% but there was no association between GY and PH amongst cultivars under either SA or IR conditions. Harvest index showed a positive association with GY under IR (R^2^=0.39, P<0.001) and SA (R^2^=0.28, P<0.001) conditions (Fig. 2 b).

### 3.2 Root system traits and correlations with yield and yield components

In 2018, root traits were not significantly affected by the irrigation treatment. However, differences between the genotypes were observed in all the root traits (P<0.05, Table 3). Overall root surface area ranged from 28.4 to 59.4 cm^2^ per plant with mean of 44.3 cm^2^ per plant and 22.9 to 60.1 cm^2^ per plant with mean of 43.4 cm^2^ per plant under IR and SA conditions respectively. Interestingly under SA conditions, GY and AGDM showed negative association with root surface area (r = −0.29 and r = −0.32), and root volume (r = −0.26 and r = −0.28). However, TGW showed positive association with root surface area (r = 0.40), root diameter (r = 0.52), and root volume (r = 0.49). Under IR conditions root diameter and TGW showed positive association with GY (r = 0.27, and r = 0.29 respectively). Also, under IR conditions onset of senescence showed a positive association with root diameter (r = 0.29, Table 5).

**Table 3.** ANOVA showing significance for genotype (G), treatment (T), interaction (G x T) and genetic ranges for root and senescence traits: root surface area (RoSuAr), root diameter (RoDiM), root volume (RoVol), NDVI at anthesis (NDVI), senescence start (SenSt), senescence duration (SenDu) in 2018.

|  | RoSuAr (cm<sup>2</sup>) |  | RoDiM (mm) |  | RoVol (cm<sup>3</sup>) |  | NDVI |  | SenSt (°CD) |  | SenDu (°CD) |  |
| --- | --- | --- | --- | --- | --- | --- | --- | --- | --- | --- | --- | --- |
|  | IR | SA | IR | SA | IR | SA | IR | SA | IR | SA | IR | SA |
| Mean | 44.3 | 43.4 | 0.52 | 0.51 | 0.57 | 0.56 | 0.63 | 0.52 | 648 | 250 | 1001 | 1501 |
| Min | 28.4 | 22.9 | 0.42 | 0.41 | 0.31 | 0.24 | 0.56 | 0.40 | 425 | 149 | 669 | 1131 |
| Max | 59.4 | 60.1 | 0.69 | 0.64 | 0.86 | 0.87 | 0.71 | 0.61 | 1033 | 461 | 1401 | 2221 |
| %red |  | 2.17 |  | 1.36 |  | 2.82 |  | 17.6 |  | 61.4 |  | -49.9 |
| **LSD** |  |  |  |  |  |  |  |  |  |  |  |  |
| G | 11.6*** |  | 0.08*** |  | 0.19*** |  | 0.56*** |  | 127*** |  | 243*** |  |
| T | 32.6 |  | 0.25 |  | 0.13 |  | 1.45 T |  | 114** |  | 8.43*** |  |
| T x G | 17.13 |  | 0.12 |  | 0.26 |  | 0.82* |  | 179*** |  | 341*** |  |

Significance levels displayed as ns>.10, T <.10 & > .05, \* <.05 > .01, \*\* <.01, \*\*\*<0.001.

In 2019, root diameter (RoDiM), root number per plant (RoNoPl) and root:shoot ratio (Ro:Sh ratio) showed significant differences between genotypes and treatments (Table 4). Overall phenotypic variation amongst genotypes for root angle was from 46.7° to 68.0° with mean of 56.6° and 46.1° to 63.8° with mean of 56.3° under IR and SA conditions, respectively. Root number per shoot (RoNoSh) showed a positive association with GY and HI (r = 0.32 and r = 0.35, respectively) under SA conditions but there was no association under IR conditions (Table 5). AGDM showed a negative association with root diameter (r = −0.29) and root dry weight per plant (r = −0.49) under IR conditions. Wider root angle was also associated with higher GY and AGDM under IR conditions (r = 0.29 and r = 0.33, respectively; Table 6). Narrower root angle was associated with more roots per plant under SA conditions, whereas under IR conditions these associations were not significant. Root dry weight per plant also showed a positive association with root diameter under IR conditions. There was a strong positive association between root dry weight and root number per plant under both SA and IR conditions.

**Table 4.** ANOVA showing significance for genotype (G), treatment (T), interaction (G x T) and genetic ranges for root and senescence traits: root angle (RoAng), root diameter (RoDiM), root dry weight per plant (RoDrWtPl), root number per plants (RoNoPl), root:shoot ratio (Ro:Sh Ratio), canopy temperature (CT), canopy green area per meter square at anthesis (GA) and NDVI at anthesis (NDVI) studied in 2019.

|  | RoAng (°) |  | RoDiM (mm) |  | RoDrWtPl (g) |  | RoNoPl |  | Ro:Sh Ratio |  | CT (°C) |  | GA |  | NDVI |  |
| --- | --- | --- | --- | --- | --- | --- | --- | --- | --- | --- | --- | --- | --- | --- | --- | --- |
|  | IR | SA | IR | SA | IR | SA | IR | SA | IR | SA | IR | SA | IR | SA | IR | SA |
| Mean | 56.6 | 56.3 | 0.71 | 0.68 | 0.52 | 0.49 | 19.9 | 15.2 | 0.08 | 0.06 | 29.3 | 36.9 | 1.94 | 1.68 | 0.72 | 0.57 |
| Min | 46.7 | 46.1 | 0.46 | 0.50 | 0.15 | 0.20 | 13.8 | 10.8 | 0.03 | 0.02 | 26.0 | 31.5 | 1.82 | 1.43 | 0.64 | 0.46 |
| Max | 68.0 | 63.8 | 0.87 | 0.79 | 0.99 | 0.99 | 27.0 | 23.3 | 0.17 | 0.10 | 32.0 | 43.0 | 2.00 | 1.90 | 0.79 | 0.66 |
| %red |  | 0.51 |  | 4.54 |  | 6.62 |  | 23.9 |  | 27.6 |  | -26.2 |  | 13.1 |  | 20.8 |
| **LSD** |  |  |  |  |  |  |  |  |  |  |  |  |  |  |  |  |
| G | 9.09 |  | 0.13* |  | 0.32 |  | 5.28* |  | 0.04 T |  | 4.19 |  | 0.13** |  | 0.06*** |  |
| T | 1.82 |  | 0.03** |  | 0.06 |  | 1.06*** |  | 0.01*** |  | 0.84*** |  | 0.02*** |  | 0.01*** |  |
| T X G | 12.9 |  | 0.18 |  | 0.45 |  | 7.47 |  | 0.05 |  | 5.93 |  | 0.19 |  | 0.08* |  |

Significance levels displayed as ns>.10, T <.10 & > .05, \* <.05 > .01, \*\* <.01, \*\*\*<0.001.

**Table 5.** Correlation matrix showing correlation coefficient (r) values for grain yield (GY), above ground dry matter (AGDM), harvest index (HI), thousand grain weight (TGW), heading date (HD), root surface area (RoSuAr), root diameter (RoDiM), root volume (RoVol), NDVI at anthesis (NDVI), NDVI senescence start (SenSt), NDVI senescence duration (SenDu). Below diagonal IR and above diagonal SA for 2018.

|  | GY | AGDM | HI | TGW | HD | RoSuAr | RoDiM | RoVol | NDVI | SenSt | SenDu |
| --- | --- | --- | --- | --- | --- | --- | --- | --- | --- | --- | --- |
| GY | -- | 0.87 *** | 0.51 *** | 0.04 | -0.35 ** | -0.29 * | -0.08 | -0.26 T | 0.34 ** | 0.14 | -0.10 |
| AGDM | 0.72 *** | -- | 0.03 | -0.02 | -0.30 * | -0.32 * | -0.10 | -0.28 * | 0.23 | 0.03 | 0.02 |
| HI | 0.21 | -0.51 *** | -- | 0.09 | -0.25 T | -0.05 | 0.00 | -0.05 | 0.27 T | 0.26 T | -0.25 T |
| TGW | 0.29 * | -0.01 | 0.40 ** | -- | -0.18 | 0.40 ** | 0.52 *** | 0.49 *** | 0.01 | 0.01 | 0.07 |
| HD | -0.11 | 0.18 | -0.38 ** | -0.28 * | -- | 0.20 | -0.14 | 0.12 | 0.10 | -0.14 | -0.14 |
| RoSuAr | 0.02 | 0.03 | 0.02 | 0.23 | 0.31 * | -- | 0.37 ** | 0.91 *** | -0.06 | -0.07 | 0.11 |
| RoDi | 0.27 * | 0.11 | 0.16 | 0.43 *** | -0.23 | 0.16 | -- | 0.68 *** | -0.12 | 0.03 | 0.04 |
| RoVo | 0.16 | 0.09 | 0.10 | 0.43 ** | 0.13 | 0.84 *** | 0.65 *** | -- | -0.09 | -0.03 | 0.09 |
| NDVI | 0.34 ** | 0.36 ** | -0.07 | -0.03 | 0.26 T | -0.03 | -0.14 | -0.08 | -- | 0.48 *** | -0.63 *** |
| SenSt | 0.38 ** | 0.30 * | 0.03 | 0.35 ** | -0.33 * | 0.08 | 0.29 * | 0.20 | -0.16 | -- | -0.76 *** |
| SenDu | -0.27 * | -0.17 | -0.05 | -0.25 T | 0.11 | -0.16 | -0.26 T | -0.25 T | 0.13 | -0.74 *** | -- |

Significance levels displayed as ns>.10, <.10 T> .05, \* <.05>.01, \*\* <.01, \*\*\*<0.001.

**Table 6.**
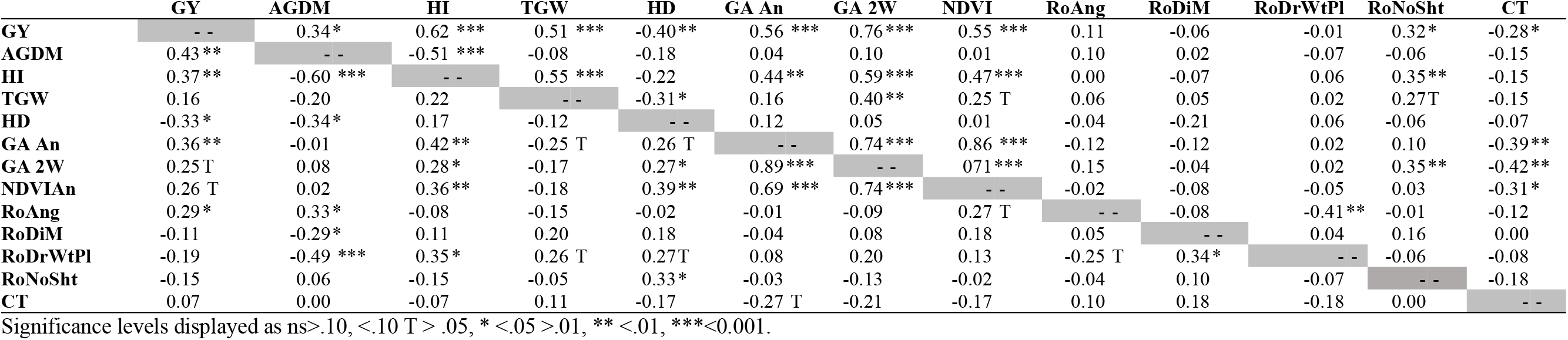
Correlation matrix showing correlation coefficient (r) values for grain yield (GY), above ground dry matter (AGDM), harvest index (HI), thousand grain weight (TGW), days to heading (DH), canopy green area per meter square at anthesis (GA An) and after 2 weeks of anthesis (GA 2W), NDVI at anthesis (NDVI), root angle (RoAng), root diameter (RoDiM), root dry weight per plant (RoDrWtPl), root number per shoot (RoNoSht), and canopy temperature at anthesis (CT). Below diagonal IR and above diagonal SA for 2019.

|  | GY | AGDM | HI | TGW | HD | GA An | GA 2W | NDVI | RoAng | RoDiM | RoDrWtPl | RoNoSht | CT |
| --- | --- | --- | --- | --- | --- | --- | --- | --- | --- | --- | --- | --- | --- |
| GY | -- | 0.34 * | 0.62 *** | 0.51 *** | -0.40 ** | 0.56 *** | 0.76 *** | 0.55 *** | 0.11 | -0.06 | -0.01 | 0.32 * | -0.28 * |
| AGDM | 0.43 ** | -- | -0.51 *** | -0.08 | -0.18 | 0.04 | 0.10 | 0.01 | 0.10 | 0.02 | -0.07 | -0.06 | -0.15 |
| HI | 0.37 ** | -0.60 *** | -- | 0.55 *** | -0.22 | 0.44 ** | 0.59 *** | 0.47 *** | 0.00 | -0.07 | 0.06 | 0.35 ** | -0.15 |
| TGW | 0.16 | -0.20 | 0.22 | -- | -0.31 * | 0.16 | 0.40 ** | 0.25 T | 0.06 | 0.05 | 0.02 | 0.27 T | -0.15 |
| HD | -0.33 * | -0.34 * | 0.17 | -0.12 | -- | 0.12 | 0.05 | 0.01 | -0.04 | -0.21 | 0.06 | -0.06 | -0.07 |
| GA An | 0.36 ** | -0.01 | 0.42 ** | -0.25 T | 0.26 T | -- | 0.74 *** | 0.86 *** | -0.12 | -0.12 | 0.02 | 0.10 | -0.39 ** |
| GA 2W | 0.25 T | 0.08 | 0.28 * | -0.17 | 0.27 * | 0.89 *** | -- | 0.71 *** | 0.15 | -0.04 | 0.02 | 0.35 ** | -0.42 ** |
| NDVIAn | 0.26 T | 0.02 | 0.36 ** | -0.18 | 0.39 ** | 0.69 *** | 0.74 *** | -- | -0.02 | -0.08 | -0.05 | 0.03 | -0.31 * |
| RoAng | 0.29 * | 0.33 * | -0.08 | -0.15 | -0.02 | -0.01 | -0.09 | 0.27 T | -- | -0.08 | -0.41 ** | -0.01 | -0.12 |
| RoDiM | -0.11 | -0.29 * | 0.11 | 0.20 | 0.18 | -0.04 | 0.08 | 0.18 | 0.05 | -- | 0.04 | 0.16 | 0.00 |
| RoDrWtPl | -0.19 | -0.49 *** | 0.35 * | 0.26 T | 0.27 T | 0.08 | 0.20 | 0.13 | -0.25 T | 0.34 * | -- | -0.06 | -0.08 |
| RoNoSht | -0.15 | 0.06 | -0.15 | -0.05 | 0.33 * | -0.03 | -0.13 | -0.02 | -0.04 | 0.10 | -0.07 | -- | -0.18 |
| CT | 0.07 | 0.00 | -0.07 | 0.11 | -0.17 | -0.27 T | -0.21 | -0.17 | 0.10 | 0.18 | -0.18 | 0.00 | -- |

Significance levels displayed as ns>.10, <.10 T> .05, \* <.05>.01, \*\* <.01, \*\*\*<0.001.

### 3.3 Canopy senescence and temperature traits

Overall, in 2018 NDVI at anthesis (NDVI, GS61) ranged from 0.56 to 0.71 and 0.40 to 0.61 under IR and SA conditions, respectively. ANOVA shows that there was a significant difference between genotypes and irrigation treatments along with significant G x T interaction for NDVI at anthesis, senescence start (SenSt) and senescence duration (SenDu) (Table 3). There was a positive association between NDVI at anthesis and GY under both, IR (r = 0.34) and SA (r = 0.34) conditions (Table 5). Under IR conditions, NDVI at anthesis and senescence start (SenSt) showed positive associations with GY (r = 0.34 and r = 0.38, respectively) whereas senescence duration (SenDu) showed negative association with GY (r = −0.27) (Table 5).

In 2019, NDVI at anthesis ranged from 0.64 to 0.79 with mean of 0.72 and 0.46 to 0.66 with mean of 0.57 under IR and SA conditions, respectively. There was significant difference between genotypes and treatments for canopy green area per meter square at anthesis and NDVI at anthesis. G x T interaction was significant only for NDVI at anthesis (Table 4). In terms of association between GY and vegetation indexes, canopy green area per meter square at anthesis showed better association with GY than NDVI at anthesis and these associations were stronger under SA (r = 0.56 and r = 0.55, respectively) conditions than IR (r = 0.36 and r = 0.26, respectively) conditions (Table 6, Fig. 3). Under SA conditions both canopy green area at anthesis and NDVI at anthesis showed a negative correlation (r = −0.39 and r = −0.31, respectively) with canopy temperature (Table 6; Fig. 3). Canopy green area and NDVI after two and three weeks of anthesis was not associated with GY.

**Figure 3:**
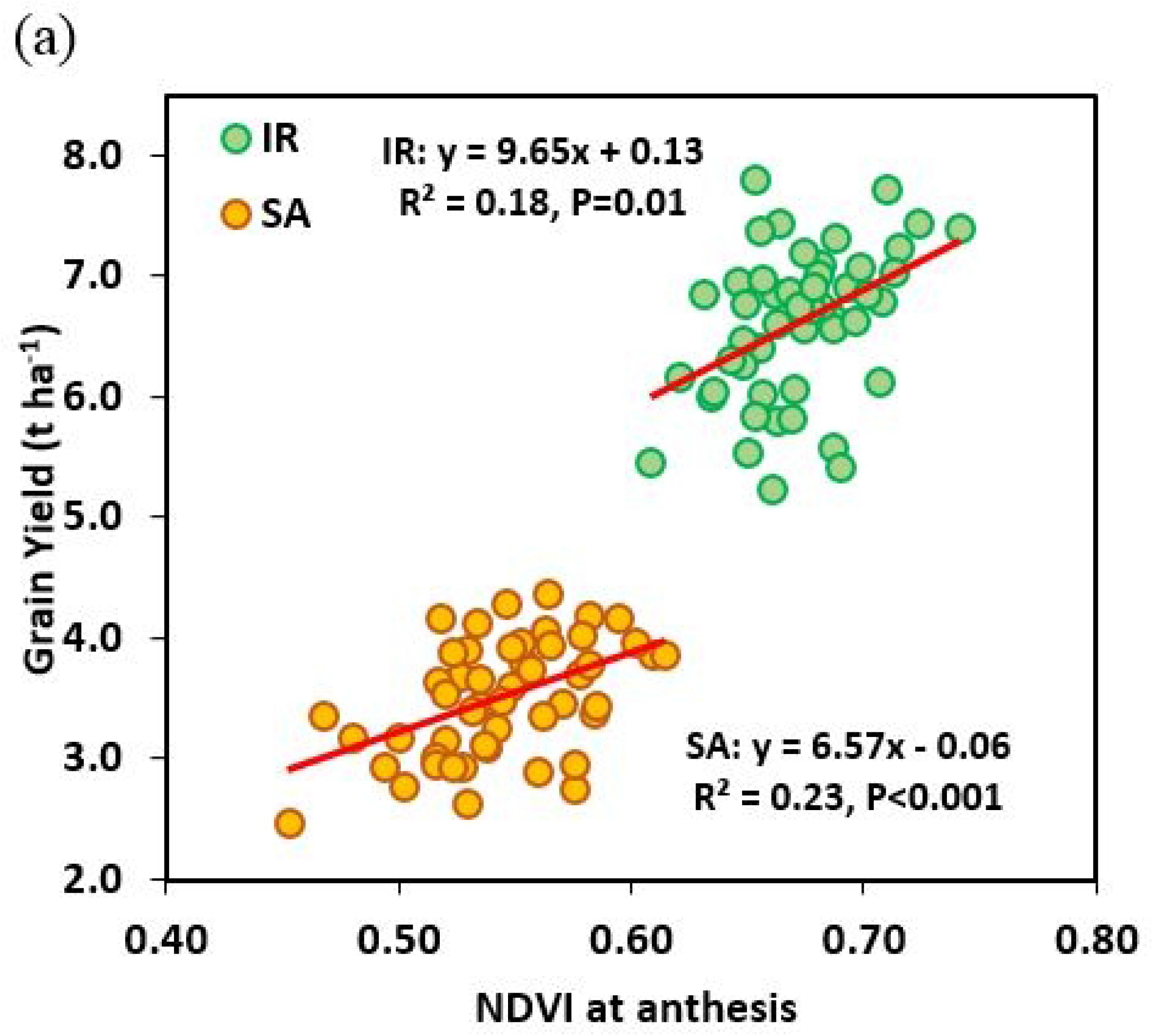
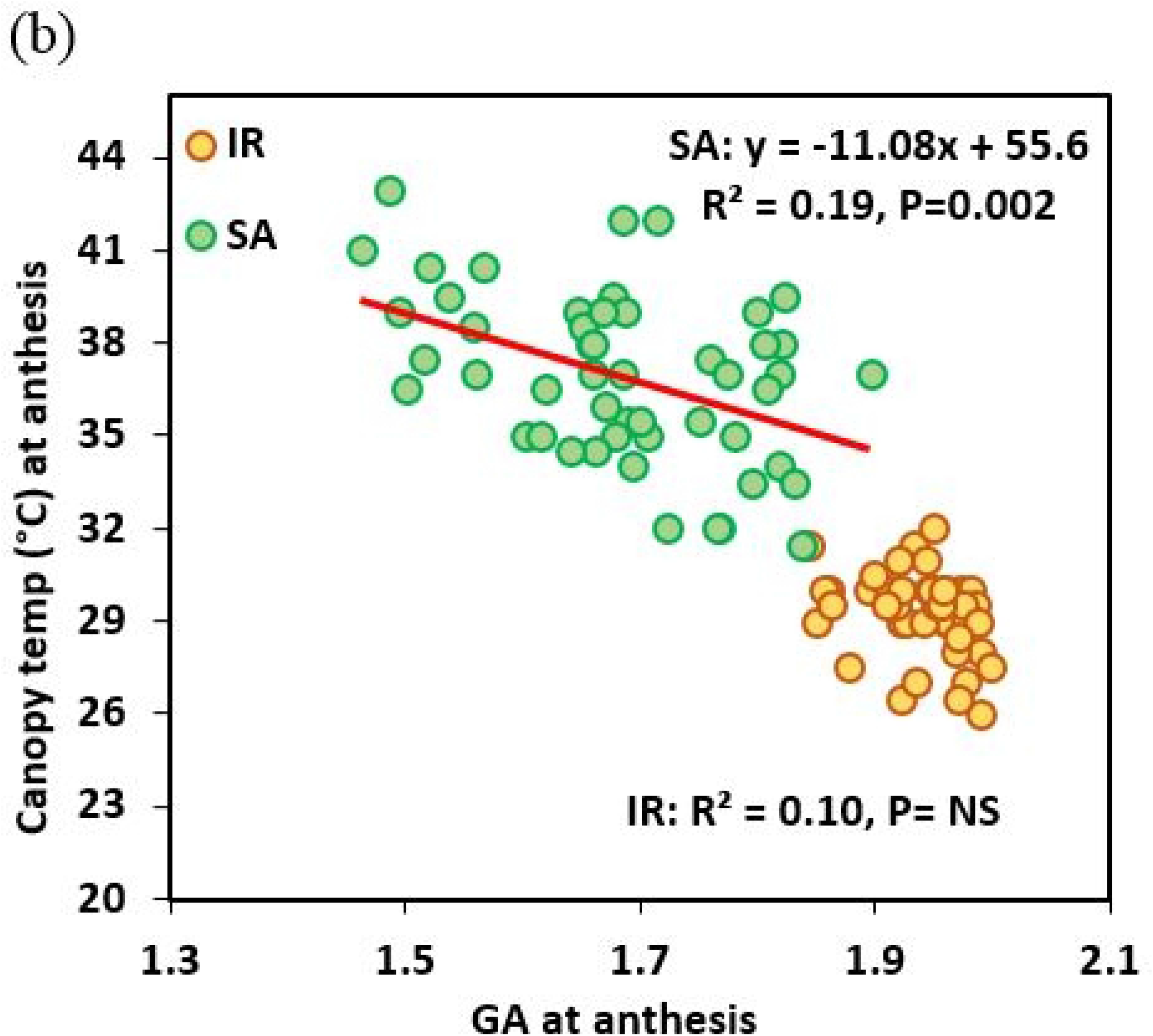
Linear regression amongst 50 wheat genotypes between GY and (a) NDVI at anthesis (b) Green area per meter square (GA) at anthesis under irrigated (IR) and semiarid (SA) conditions (mean of 2018 and 2019) at anthesis in 2019.

### 3.4 Association between different yield components under IR and SA condition

Under IR conditions, the first principal component (PC1) explained 25.3% variation and the group of traits explaining this variation were fruiting efficiency, spikelet’s per ear, days to heading with positive effect whereas TGW showed negative effect. The second principal component (PC2) explained 20.8% variation and the group of traits explaining this variation included grains per ear, harvest index, grain yield, and root diameter with positive effects whereas above ground dry matter showed negative effects.

Under SA conditions, PC1 explained 25.6% variation and trait associated with positive effect - were grains per ear, harvest index, grain yield, and NDVI whereas days to heading had negative effects. PC2 explained 21.0% of variation and traits showing association with positive effects were - thousand grain weight, plant height, root diameter, whereas spikelet’s per ear had negative effect (Fig. 4).

**Figure. 4:**
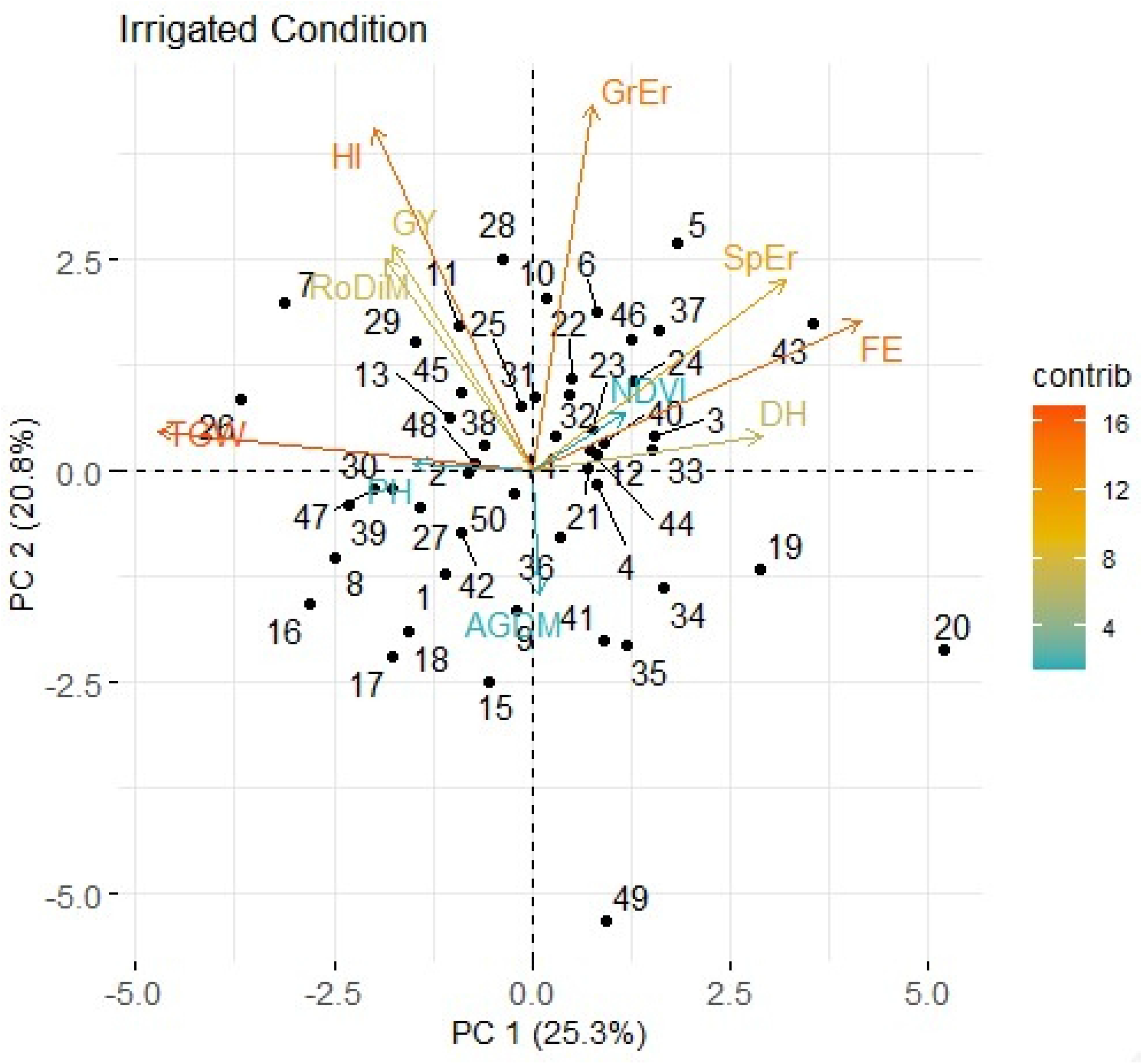
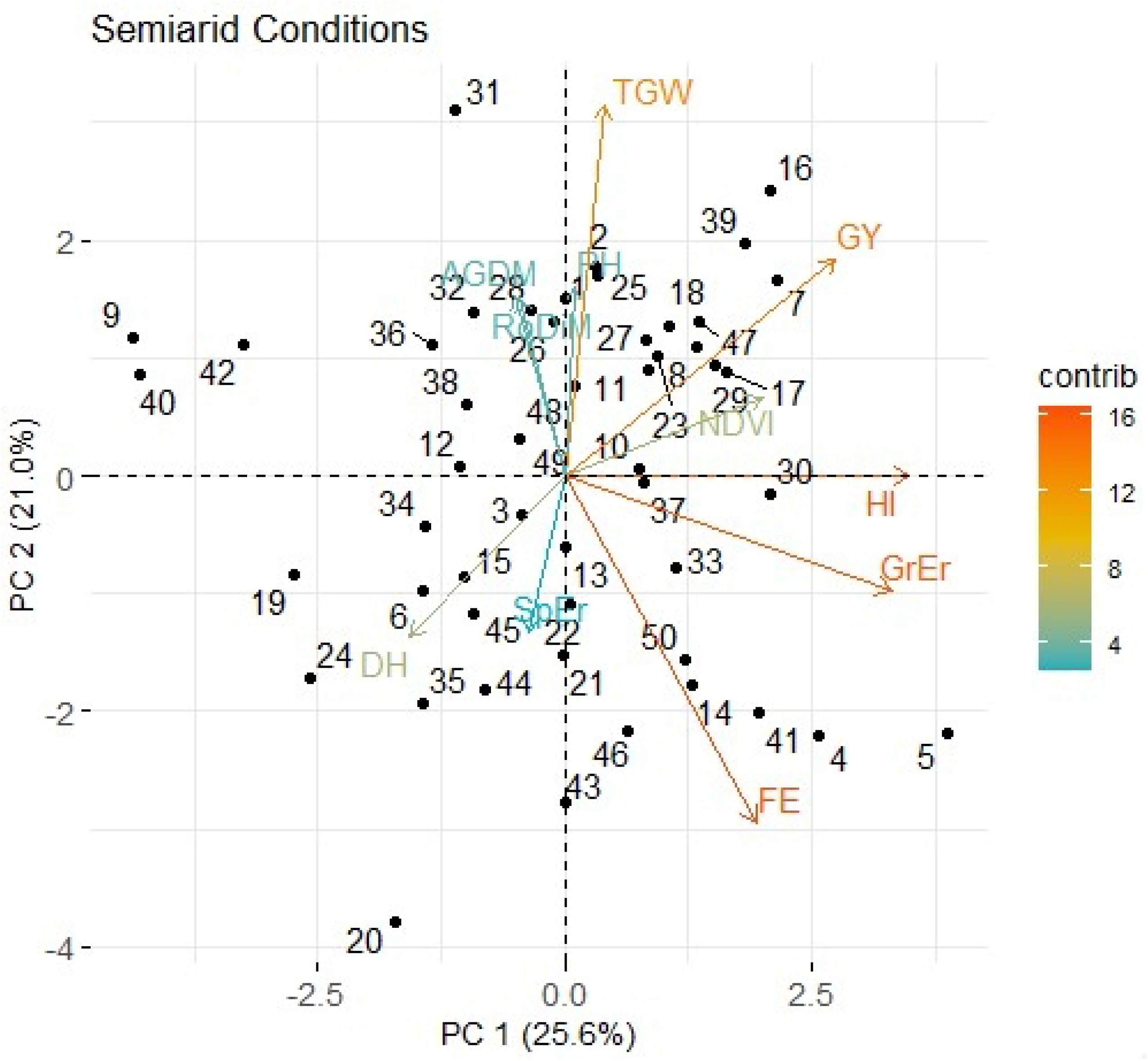
Bi-plot for grain yield (GY), above ground dry matter (AGDM), harvest index (HI), thousand grain weight (TGW), days to heading (DH), grain numbers per ear (GrEr), spikelet number per ear (SpEr), fruiting efficiency at harvest (FE), NDVI at anthesis (NDVI) and root diameter (RoDiM) under irrigated and semiarid conditions for 50 cultivars (Mean of 2018 and 2019).(Contrib: contribution in total variation in per cent – red to blue: stronger to low).

### 3.5 Marker-traits associations

Marker, allele/haplotype effect, mean of traits for each allele and probability of significant difference in the mean under IR and SA conditions are presented in Table 7 for 2018 and 2019. In 2018, marker-trait analysis showed that marker *TaCwi-4A* have significant influence on GY under SA. This marker was responsible for increase in GY by 17.1% under SA in presence of Hap-4A-C allele which is responsible for higher yield under drought conditions (32) and present in 59% of genotypes. Marker *Dreb1* was responsible for increase root surface area (RoSuAr) and root volume (RoVol) only under SA conditions. *Dreb1* allele *TaDreb-B1a* increased these traits by 9.4% and 13.3%, respectively, under SA conditions. *Dreb1* was also associated with extended SenEnd under SA conditions. The *PRR73-A1* gene had a significant influence on GY under SA but not under IR conditions. There was an increase of 14.1% under SA conditions in presence of Hap-II allele which was present in 29% of genotypes studied. This allele was also associated with extended senescence end (SenEnd) under SA and IR conditions.

**Table 7.**
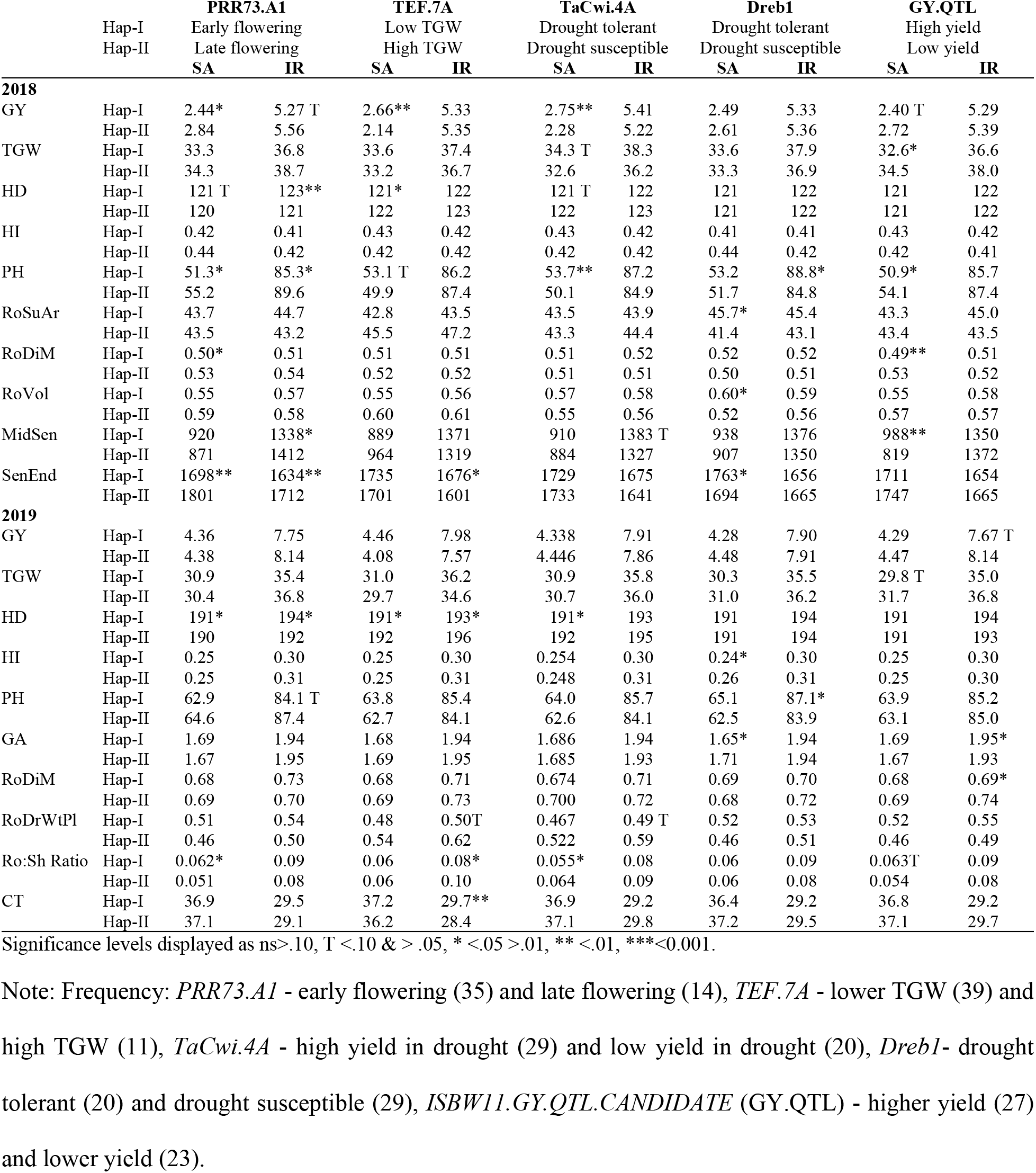
The key markers and haplotype mean for traits: grain yield (GY t ha^−1^), harvest index (HI), plant height (PH cm), thousand grain weight (TGW g), heading date (HD DAS), root surface area (RoSuAr cm^2^), root diameter (RoDiM mm), root volume (RoVol cm^3^), NDVI senescence end (SenEnd °CD), NDVI Mid senescence (MidSen °CD), canopy green area per meter square at anthesis (GA m2), root angle (RoAng °), root dry weight per plant (RoDrWtPl g), root shoot ratio (Ro:Sh ratio), and canopy temperature (CT °C) under SA and IR conditions along with significant difference between the two haplotype mean to show marker-trait association in 2018 and 2019 for 50 genotypes.

The marker-trait associations in 2018 were not apparent for the same traits in 2019. However, Marker *PRR73-A1* with allele Hap-I (early flowering) was responsible for increase above ground biomass at anthesis (AGBMAn; data not shown) and root: shoot ratio (Ro:Sh Ratio).

## 4. Discussion

### 4.1 Grain yield responses to drought and association with phenology

In our experiments, there was a moderately severe drought with yield reducing overall by 3.06 t ha^−1^ (−47%). This is representative of Mediterranean drought effects in semi-arid conditions reported for wheat with reductions of yield typically ca. 30-50% (39). The cultivars responded differently to the drought stress as indicated by the significant irrigation x genotype interaction. Higher yield under IR conditions was associated with greater yield loss under SA conditions amongst the cultivars. From the physiological standpoint, it is not surprising that absolute reduction in yield for a given reduction in water resource is strongly influenced by yield potential (40–42).

Drought had only a small effect on heading date overall, advancing GS59 on average by one day in 2018 and three days in 2019; genotypes responded similarly to drought (Fig. 2). In both SA and IR conditions, HD ranged by 10 days amongst cultivars. Correlations between heading date and grain yield were negatively associated under SA conditions in both years, and there was also a negative association under IR conditions in 2019 although less strong than under SA conditions. Bi-plots for the cross-year mean also confirm these effects. Early flowering has been associated with drought escape in wheat in environments subjected to severe early season drought stress, e.g., in northern Mexico (40). Similarly Worland et al. (43) reported increased yield for *Ppd-D1a* early-flowering NILs by ca. 5% compared to *Ppd-D1b* controls in dry years. Each day’s advancement in HD raised yield by 0.11 t ha^−1^. Soil depth was more than 2 m at the field site with a very low organic matter. Presumably a shorter pre-anthesis phase reduced water uptake in this phase, so that season-long water uptake was redistributed more favorably with regard to the post-anthesis period. However, in one year a similar negative association between HD and yield was recorded under irrigated conditions. This indicated that the negative association may have been partly associated with advanced flowering leading to cooler prevailing temperatures during grain filling and therefore more calendar days for grain filling (44). (45) showed that during grain filling, an increase in 1°C mean daily temperature higher than optimum can be responsible for decrease in ca. 2.8 mg of grain weight. Observations regarding flowering time and drought resistance are very much dependent on the exact timing of drought stress and we recognize that the present experiments would need to be repeated over more years before we could conclude with certainty that later heading date overall has a negative effect on yield losses under droughts in Turkey. For example, it may be that there is a trade-off between early flowering and the development of a smaller root system, as suggested by (46).

### 4.2 Associations between canopy senescence traits and responses to drought

In the present study, greater yield amongst cultivars was associated with higher green area per meter square and NDVI at around heading or anthesis under both drought and irrigated conditions. This likely reflected an association between NDVI and biomass at anthesis and hence grains m^−2^ under both treatments. Genetic variation in GY in wheat has previously been associated with green canopy area duration under drought in wheat (41,47–50), sorghum (51). The role of senescence dynamics - start, end and rate of senescence - is important as they relate to grain filling duration and post-anthesis water and N uptake (50,52). Under irrigated conditions, GY was associated positively with senescence start (SenSt) in 2018. Genotypes having delayed senescence start later may be able to accumulate more plant nutrients and carbohydrates during grain filing duration resulting in higher yield. Our results suggested that there was source limitation if grain growth even under sufficient soil moisture. This could have been due to some heat stress incurred in the experiments combined with the higher grain number in irrigated conditions leading to source limitation during the later stages of grain filling (53).

One of the objectives of this experiment was to compare the two, NDVI and RGB-based vegetation indexes as methods for measuring canopy green area. We found that the RGB-based vegetation indexes showed better association with GY than NDVI measured with the handheld Trimble GreenSeeker (Trimble Navigation Ltd, USA) and in addition it increase throughput. Better association of grain yield with RGB based vegetation indexes than NDVI was also reported in durum wheat by (26) and (27). The role of cooler canopy at anthesis associated with water uptake and root system for drought tolerance is previously reported (5) and was also observed in this experiment with a negative association between GA at anthesis, NDVI at anthesis and GY with canopy temperature (CT) under SA conditions in 2019.

### 4.3 Effect of shovelomics root traits on responses to drought

Shovelomics represents a high-throughput phenotyping method for field-grown crops and has been used to quantify genetic variation in root traits in maize (18–20), legumes (54) and barley (21). (22) carried out field shovelomics for durum wheat recombinant inbred lines (RIL) for crown root length, number and angle and reported QTL. In our study we applied a shovelomics methodology for bread wheat to quantify variation in nodal root angle, length, roots plant^−1^ and roots shoot^−1^ and association with GY. In the present study, the range in nodal root angle of 45-65° in the semiarid treatment was similar to that of 42.3-69.2° reported by (22) for the Colosseo × Lloyd durum wheat mapping population assessed at anthesis in the field under optimum agronomic conditions in Italy. Values under irrigation of 45.0-70.0° were also similar to those reported by (22) under irrigation. The association of shallower root angle with higher GY under irrigated conditions in 2019 was possibly associated with increased root density at shallower depth increased recovery of fertilizer N uptake which is predominately distributed in the top 30 cm of the soil. Shallower roots may also increase uptake of irrigation water leading to more transpiration and cooler canopies, so avoiding heat stress. Generally shallower root angle is associated with shallower root distribution in soil and narrow root angle associated with deeper root distribution in soil (15). In durum wheat, (55) reported 20 to 40% yield advantage under irrigated conditions with the shallowest root types compared to deep root type.

(56) reported 46.2% and 68.3% of wheat roots are distributed in upper 15 and 30 cm, respectively. In contrast, a steeper angle would be expected to correlate with relatively deeper roots and greater yield under SA conditions, as has been reported in wheat in Australia (13–15). In our experiments soil depth was more than 2 m. Previous studies in maize found steeper root angle related to increased rooting depth under low nitrogen field environments in the USA and South Africa (18). However, in our results we did not see a significant association between root angle and GY under drought. However, under SA conditions in 2019 there was a positive association between nodal root number per shoot and GY and HI, but no association under IR conditions. These results suggest that the wheat ideotype for drought tolerance may be a plant with relatively few tillers but a high number of nodal roots per shoot associated with steeper roots and increased rooting depth. A field study in Pennsylvania in maize found that reduced nodal roots per plant led to increased root length at depth and 57% higher grain yield under water-stressed conditions (57). Tillering influences carbon partitioning, and there is some evidence that reduced tillering increases rooting depth in wheat (58,59) and rice (60).

In 2018 terminal stress was severe as rainfall was lower towards the end of crop season than in 2019 (Table 1). In 2018 under semiarid conditions genotypes which had less root surface area and root volume per plant showed higher yield and biomass. It could be speculated that less surface roots may have been associated with more roots distributed relatively deeper in 2018 under SA conditions. Overall under drought conditions fine roots with smaller root diameter may have advantage in exploring maximum area in soil with less energy invested to grow them (61). Whereas in 2019, when terminal drought was less severe, this association with root surface area and volume was not observed.

Shovelomics is becoming an increasingly popular method for the high-throughput phenotyping of field-grown crop roots. The shovelomics method we have applied for phenotyping nodal root traits in winter wheat in the present study was shown to be a valuable technique. However, association of canopy temperature with GY indicates that water uptake at depth was contributing to yield increase. As shovelomics traits measures root traits in top 30 cm of soil profile, these technique requires further validation that it is a reliable indicator for roots at depth. Shovelomics is a relatively high-throughput method, making it possible to sample, wash and measure root crowns from 200 plots in two person-days. Shovelomics phenotyping platform is significantly faster the field soil coring, which would take approximately one person-month to sample, wash and extract roots, and image the samples for 200 plots in the present study. This high-throughput shovelomics platform can potentially be used to phenotype large populations to identify QTL, search for candidate genes and develop molecular marker for marker-assisted selection. There are examples of deploying QTL for root depth in other cereal species. In rice, the *Dro1* gene related to steeper crown root angles and deeper rooting was identified by measuring nodal root traits in a high-throughput controlled environment study (62) and has since been used to produce drought tolerant NILs which have been phenotyped in field conditions (63).

### 4.4 Association between molecular markers and responses to drought

Genomic studies using high-throughput genotyping assays like KASP have made it possible to genotype large populations at various loci within a very short time (38). Several recent studies used KASP markers to identify the allelic variation of functional genes in wheat cultivars from China (38), United States (64), and Canada (65). In our study the clear association of *TEF-7A* and *TaCwi-4A* with GY and *Dreb1* with root surface area (RoSuAr) under SA conditions indicated the usefulness of deploying these markers in wheat breeding for drought tolerance.

Dehydration responsive element binding proteins, *Dreb1*, have been induced by water stress, low temperature and salinity (66). In this study, *TaDreb1* was associated with increased root surface area, root volume and delayed end of senescence which indicated multi-trait effects of this transcription factor which were not previously reported. The *TaCwi-4A* marker was associated with GY, root shot ratio (Ro:Sh ratio) and PH under SA conditions. It was previously reported that storage carbohydrate accumulation in drought susceptible and tolerant cultivars depends on the expression of gene for cell wall invertase (*TaCwi*) in anthers (67). The effect of drought on pollen fertility is irreversible and may cause grain loss or yield reduction under drought conditions. Since these genes tightly control sink strength and carbohydrate supply, therefore deployment of favorable alleles of these genes could maintain pollen fertility and grain number in wheat. The drought tolerance and association of yield-related traits in CIMMYT winter wheat germplasm was strongly associated with the presence of the favorable allele of *TaCwi-4A* which ultimately increased the grain sink size during drought stress. The presence of favorable allele for *TaCwi-4A* gene can enhance the selection accuracy of drought-tolerant germplasm in marker-assisted breeding. Similarly, the association of several important traits like SenEnd, GY, HD, and GA with the flowering time related gene *PRR73-A1* is interesting and indicated the expanded role of these genes in plant development. Previously, Zhang et al (2016), identified that *PRR73-A1* was associated with plant height and explained up to 7.5% of the total phenotyping variability in Chinese wheats. This gene also showed association with plant height in this experiment in 2018. It was previously observed that flowering time related genes are very important for wheat adaptability in target environments, and these genes are associated with several yield component traits (37). Our results provided a set of target genes which could be manipulated to further fine-tune the expression of important drought-tolerance traits.

## Conclusion

In the Mediterranean environment wheat is grown mostly under semiarid conditions and frequent drought effects wheat yield severely. The strategy of developing drought-tolerant wheat varieties depends on understanding and identifying below-ground and above-ground traits for drought tolerance together with use of marker-assisted selection. In this experiment we used high-throughput phenotyping techniques for characterizing root system architectural and canopy senescence dynamic traits. We conclude that higher number of crown roots per shoot was a key trait for yield increase under drought conditions. Use of RGB-based vegetation index to characterise the canopy green area dynamics could save time and increase precision in selection. Strong association of green area index with GY at flowering was encouraging and this index can be used as tool for early stage selection for higher GY. In this experiment we have evaluated five established functional genes as these genes were associated with different drought-tolerance traits in the field experiments. Genetic marker *TaCwi.4A*, responsible for drought tolerant was associated with higher GY in drought condition in this experiment and can be used for future breeding. Overall, We also provided new insight on effects of root phenotypes and physiological traits, for example the importance of root angle under irrigated conditions and roots per shoot under drought for increasing grain yield which could be important for developing drought-tolerant cultivars.

## Acknowledgments

CIMMYT-Turkey is supported by the CGIAR Research Program on Wheat and the Republic of Turkey Ministry of Agriculture and Forestry. CIMMYT thanks the Bill and Melinda Gates Foundation (BMGF) and the UK Department for International Development (DFID) for providing financial support through the grant OPP1133199 to AM. We also like to acknowledge Emel OZER and Adam for sowing the experiment, Jose Luis Araus Ortega and Jose Armando Fernández from the University of Barcelona for providing the training for image analysis software to author.

---

**S Fig 1.**
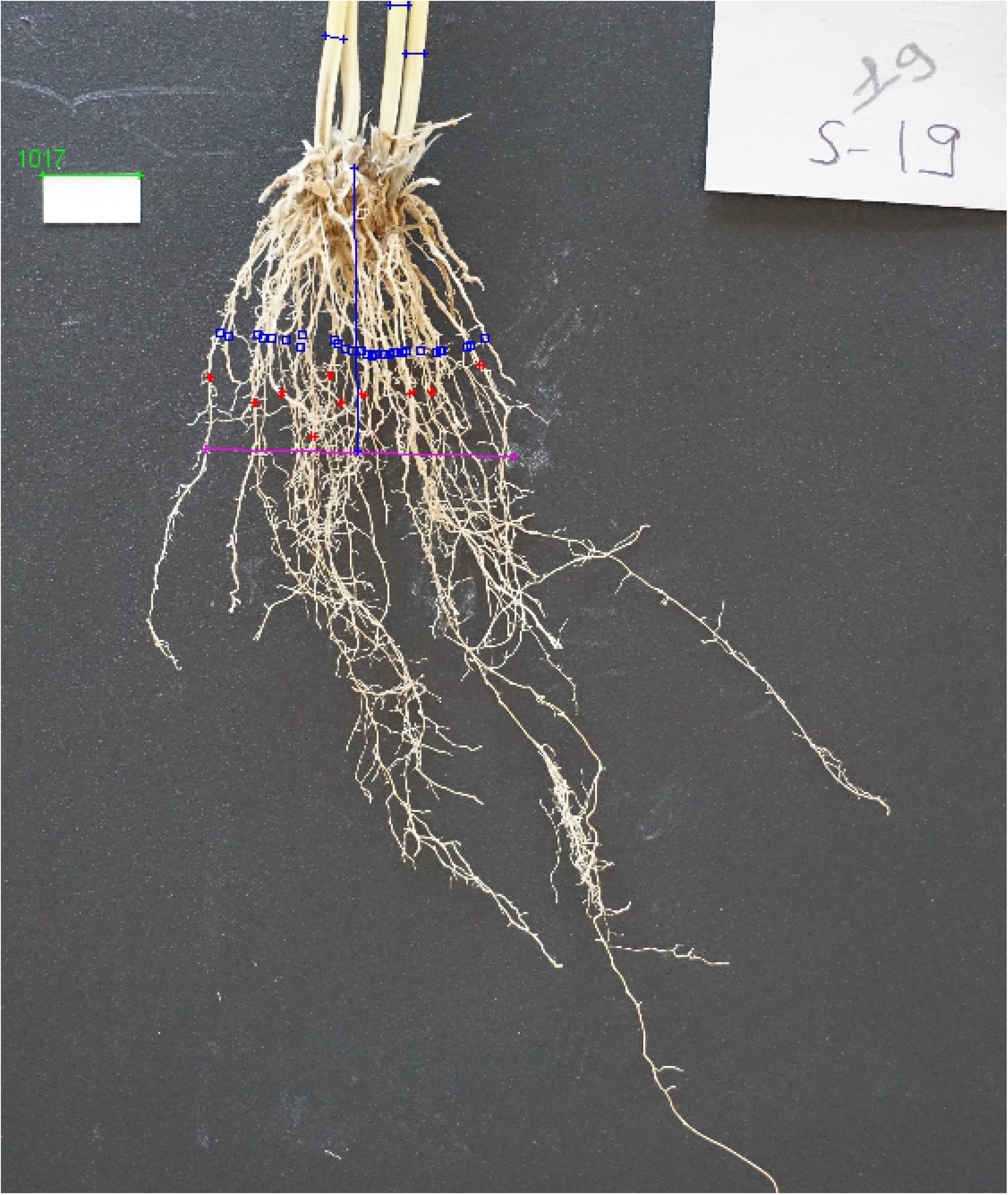
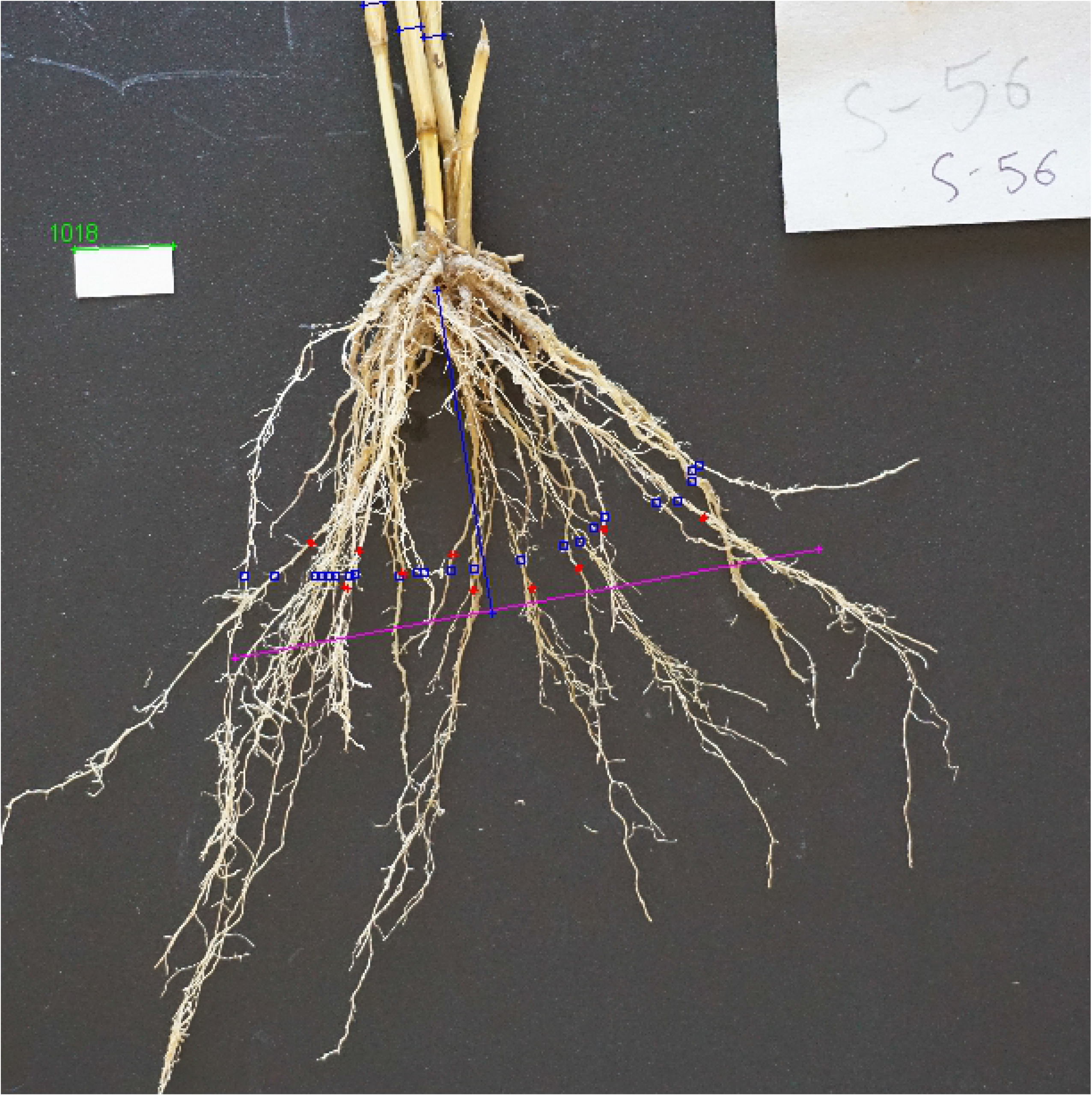

